# Driven to Extinction: On the Probability of Evolutionary Rescue from Sex-Ratio Meiotic Drive

**DOI:** 10.1101/018820

**Authors:** Robert L. Unckless, Andrew G. Clark

## Abstract

Many evolutionary processes result in sufficiently low mean fitness that they pose a risk of species extinction. Sex-ratio meiotic drive was recognized by W.D. Hamilton (1967) to pose such a risk, because as the driving sex chromosome becomes common, the opposite sex becomes rare. We expand on Hamilton’s classic model by allowing for the escape from extinction due to evolution of suppressors of X and Y drivers. We explore differences in the two systems in their probability of escape from extinction. Several novel conclusions are evident, including a) that extinction time scales approximately with the log of population size so that even large populations may go extinct quickly, b) extinction risk is driven by the relationship between female fecundity and drive strength, c) anisogamy and the fact that X and Y drive result in sex ratios skewed in opposite directions, mean systems with Y drive are much more likely to go extinct than those with X drive, and d) suppressors are most likely to become established when the strength of drive is intermediate, since weak drive leads to weak selection for suppression and strong drive leads to rapid extinction.

---

The population genetic theory of extinction was long neglected (Lewontin 1974; Orr and Unckless 2008) but has begun to receive considerable attention (Gomulkiewicz and Holt 1995; Bürger and Lynch 1997; Bell and Collins 2008; Orr and Unckless 2008; Bell and Gonzalez 2009; Gonzalez et al. 2013; Orr and Unckless 2014). This work almost exclusively deals with a population that finds itself declining due to a changed or changing *external* environment and must adapt before it becomes extinct. The population may adapt and escape extinction using either newly arising mutations or standing genetic variation, and the probabilities of these outcomes depend on several population genetic parameters.

While extrinsic environmental factors undoubtedly can threaten population persistence, *intrinsic* factors also have the potential to drive populations extinct (See Zayed and Packer 2005; Pinzone and Dyer 2013). Selfish genetic elements that skew population sex-ratios, including B chromosomes, endosymbionts and segregation distorters, may drive a population to extinction because one sex is so rare that the population cannot replace itself (Carvalho and Vaz 1999; Burt and Trivers 2006). Hamilton (1967) showed that, barring suppressors, a driving Y chromosome could cause a population of 2000 to go extinct in just 15 generations. The prognosis is only slightly improved for a driving X chromosome – extinction takes about 45 generations. In contrast, an environmental change that causes absolute fitness to decline to 0.9 will take a population of 2000 more than 70 generations to go extinct (barring a mutation that saves the population), 35 generations if fitness is 0.8 (based on calculations of Orr and Unckless 2008). After Hamilton, all theoretical work on segregation distortion has involved deterministic models with infinite population sizes (see for example Taylor and Jaenike 2002; Engelstädter *et al.* 2004; Hall 2004; Unckless and Clark 2014) except Taylor and Jaenike (2003) which dealt with the effect of sperm competition on drivers in finite populations. Of course, in many cases the parameters of the model will not lead to population extinction and a polymorphic equilibrium will be reached (Carvalho and Vaz 1999). The logical extension to Hamilton’s result has, however, been ignored: Given the invasion of a driving sex chromosome, what are the chances that a population can be rescued from extinction by a suppressor of drive either arising from new mutation or present in the standing genetic variation?

We investigate the dynamics of drive and extinction first by making some analytical progress using Hamilton’s scenario. Using the simple parameter set envisioned by Hamilton allows us to train our intuition for the more complex and more realistic cases. The models of sex ratio drive that we consider entail nonlinear recursions of several variables with ten free parameters, so numerical simulations provide a more complete picture of their behavior than the incomplete analytical solutions obtained. Of central interest is the comparison between X and Y drive and the degree to which the dynamics might provide insight regarding the empirical observation that X drive is far more common than Y drive. We find that Y drive is particularly dangerous for a population for several reasons. While the prognosis for X chromosomes is better, extinction under Y drive is likely under a wide range of parameters.

## Methods

### The Model

We examine a population of size *N* that is seeded with one driving sex chromosome at time *t*=0. In all cases males are the heterogametic (XY) sex and the driving chromosome originates in the male. While this is somewhat artificial for X drive, it saves considerable time in simulation since most single driving X chromosomes will be lost due to drift if they find themselves in females (which they would approximately two-thirds of the time if randomly assigned). Females mate *M* times at random with the available males. They then produce *R*_*0*_ offspring in total. Since this may produce more than the carrying capacity (starting population size) of offspring in the population, density-dependent factors reduce the population to that starting value every generation (unless the total population is below the starting value). This is a departure from Hamilton’s model, which allowed for population sizes to grow without limit, but is more realistic since stable populations are probably not limited solely by female productivity, but at least in part by density-dependent factors. Furthermore, with female lifetime fecundity greater than two, populations grow to unrealistic sizes very quickly without regulation. All parameters are defined in Table 1.

**Table 1.** Definition of parameters used in drive models

| Parameter | Typical range | Definition |
| --- | --- | --- |
| $N_0$ | $10^3$ to $10^6$ | Total population size before driver invades |
| $\mu$ | $10^{-8}$ to $10^{-4}$ | Rate of mutation to suppressor |
| $d$ | 0.5 to 1.0 | Strength of drive (proportion of offspring inheriting the driving sex chromosome) |
| $d_s$ | 0.5 to 0.99 | Strength of drive with suppressor (proportion of offspring inheriting the driving sex chromosome) |
| $M$ | 1 to 5 | Mean number of lifetime matings per female |
| $R_0$ | 1 to 10 | Mean number of offspring produced per female in a lifetime |
| $s_{sup\_female}$ | 0 to 0.1 | Selection coefficient for cost of suppressing X Chromosome in homozygous females (Y drive only) |
| $h$ | 0 to 1.0 | Dominance of cost of either driver (for X drive) or suppressor (for Y drive) in females |
| $s_{driver\_male}$ | 0 to 0.1 | Selection coefficient for cost of driving chromosome in males |
| $s_{sup\_male}$ | 0 to 0.1 | Selection coefficient for cost of suppressing X (in Y drive) or Y (in X drive) chromosome in males |
| $s_{driver\_female}$ | 0 to 0.1 | Selection coefficient for cost of driving chromosome in homozygous females (X drive only) |

SR males (those who possess a driving chromosome) produce *d* sperm with the sex-ratio chromosome and 1-*d* sperm with the opposite sex chromosome. For example, a male with a driving Y with *d*=1.0 produces all sons, but if *d*=0.55 he produces 55% sons. The fitness of all wildtype chromosomes is 1.0, but both driving and suppressing chromosomes may have fitness costs (see Table 1). In females, driving or suppressing X chromosomes may occur in the heterozygous or homozygous state, so a dominance value (*h*) is assigned.

Suppressors are limited to the opposite sex chromosome because a suppressor arising on the driven chromosome would simply be directly outcompeted by the unsuppressed chromosome (autosomal suppressors will be considered separately). Suppressors either arise by new mutation or are present as standing genetic variation. We treat these two possibilities separately – never allowing both new mutation and standing genetic variation in any particular realization. Suppressing chromosomes arise by new mutation from sensitive chromosomes at rate *μ*. Those present in standing genetic variation may be deleterious, neutral or nearly neutral in the absence of drive. However, instead of modeling these three cases separately, we choose an absolute number of suppressing chromosomes, and later discuss under what circumstances we might expect those starting numbers. For example, suppressors that are neutral in the absence of drive and have equal mutation rates to and from the suppressing chromosome should occur at very high (50%) frequency, while suppressors that are deleterious in the absence of drive would be found at mutation/selection balance and are therefore much more rare.

In each case there are four possible outcomes. First, the driver may be lost due to drift. Even with optimal parameters for drive this actually happens quite frequently, since the driving chromosome occurs in only one individual in generation 0. The second possible outcome is that the driving chromosome spreads and drives the population extinct as in Hamilton’s example. The third possibility is that a suppressor rises to high frequency and rescues the population from extinction. Finally, the parameters may be such that an intermediate equilibrium is reached and the population persists indefinitely even without the existence of a suppressor. This last outcome is not the focus of the current work and has been covered with deterministic models elsewhere (Carvalho and Vaz 1999).

## Simulations

We employed forward simulations that exhaustively enumerate all genotypes in the population each generation to explore the parameter space. In both the X- and Y-drive cases, the population begins with *N*/2 females with wildtype sex chromosomes, *N*/2 - 1 males with wildtype sex chromosomes and one SR male. For simulations where suppressors occur in standing genetic variation, the suppressing sex chromosomes are randomly placed in males and females (Y drive) or all in males (X drive).

Deterministic simulations were employed to address several problems such as the dynamics in Hamilton’s scenario and time to extinction without suppressors, while stochasticity was introduced for problems involving the probability of invasion (Supporting Text 1).

Stochastic simulations were conducted by grid searching input parameters and for each, iterating many replicates of the population to equilibrium.

Simulations show that when the driver is lost, the population goes extinct, the suppressor invades to a threshold (20% of the population unless otherwise noted) or a threshold number of generations has been reached (10000 unless otherwise noted). For each realization, the fate of the population is recorded as is the time to extinction (given the population went extinct), time to loss of the driver (given the driver was lost) and the time at which the suppressor arose (new mutation only). For more details on simulations, see Supplemental Text 1.

We treat the Y drive scenario first throughout because it is simpler, provides for some analytical solutions and can train the intuition for the X drive case. Though we realize that X drive is more commonly observed in natural populations, one of our objectives is to explore potential reasons for that pattern and this style of presentation facilitates such an approach.

## Results

In an attempt to simplify the *Results* section, we have placed much of the mathematical detail in Supplemental Text 2, presented results graphically when possible and set important points as numbered conclusions in italics.

### Analytical solutions using the Hamilton Model

Hamilton (Hamilton 1967) admittedly examined only the simplest of cases of his sex-ratio meiotic drive. In his model for Y drive, the starting population contains 1000 males (one with a driving Y) and 1000 females. He considered a driving Y with no fitness cost to males and perfect drive (*d*=1). Females mate once and produce two offspring. Under these simple conditions, the driving Y exterminated the population in 15 generations. Hamilton’s model for X drive was similar except that he assumed that each male could mate twice. This resulted in an initial population expansion since all females would be mated even if the sex ratio was biased toward females (until there were twice as many females as males). The driving X therefore takes about 45 generations to exterminate the population. Hamilton presented the above results graphically and verbally, but did not provide any analytical solutions. We begin by expanding on Hamilton’s model below.

#### Expansion of Hamilton’s model for Y drive

For the driving Y model, the actual number of individuals of each genotype is denoted by XX, XY and XY^SR^. As shown in Supplemental Text 2, the population size in the next generation is *N′* = 2*XX*, and iterating over several generations we find the expected population size at time *t* is

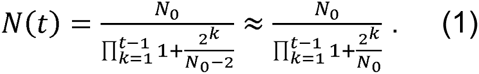

The total population size and number of each genotype are shown in figure (1A). Interestingly, Equation (1) shows that Hamilton’s population was not doomed to such rapid extinction simply because it started with a small population size (2000). Populations go extinct in 19, 23 and 27 generations for starting populations of 10^4^, 10^5^ and 10^6^ respectively, approximately linearly with the log of *N*_*0*_. This, of course, makes sense since the population size declines approximately logistically. The expected sex ratio at time t is

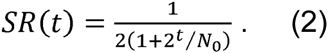

**Figure 1:**
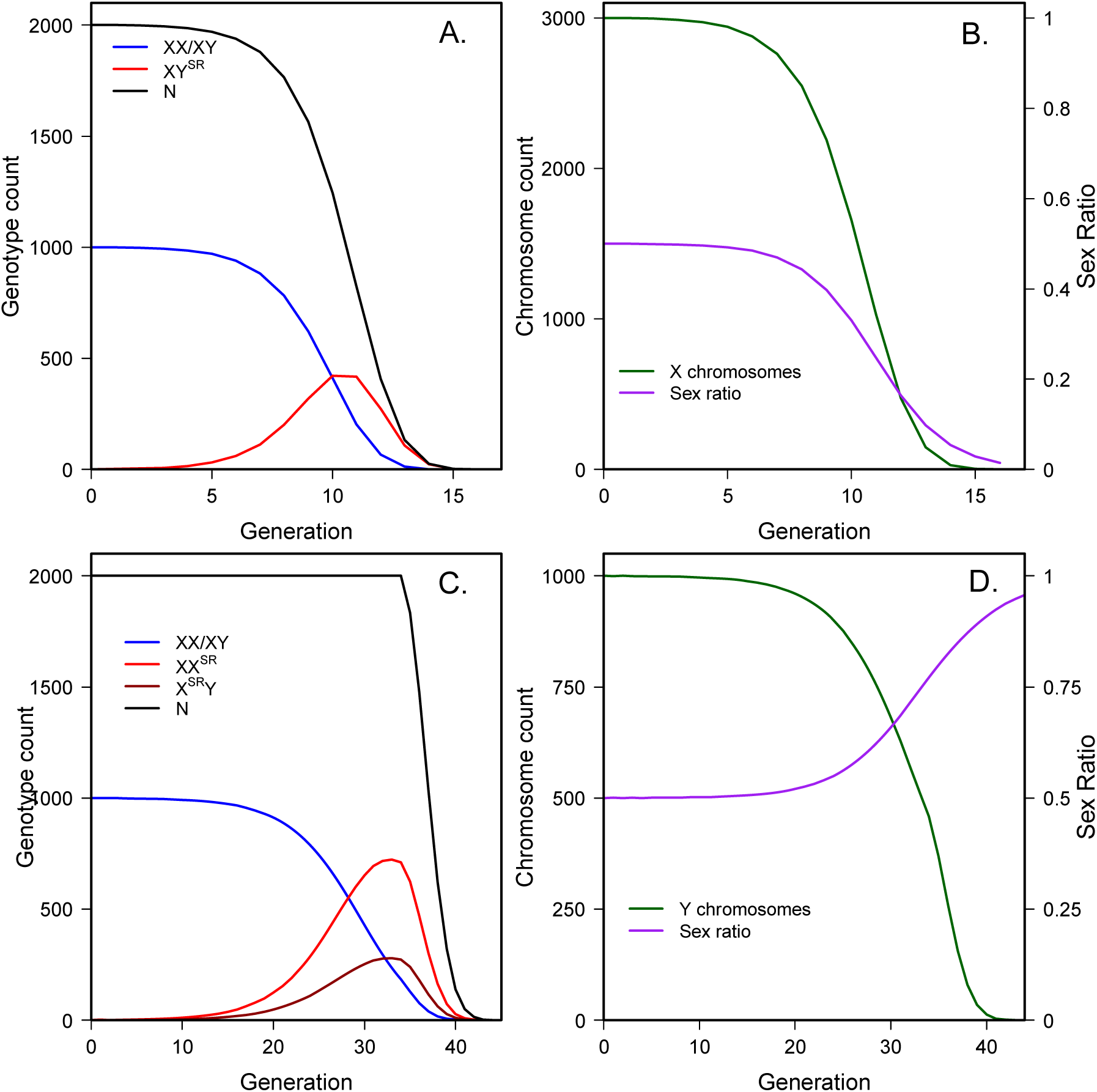
Using parameters employed by Hamilton, population extinction is rapid for both Y and X drive, A) total population size and number of each genotype under Hamilton’s Y drive scenario, B) number of mutational targets and sex-ratio under Hamilton’s Y drive scenario, C) total population size and number of each genotype under Hamilton’s X drive scenario, D) number of mutational targets and sex-ratio under Hamilton’s X drive scenario.

This illustrates why the time to extinction is not very sensitive to *N*_*0*_: a sex ratio that is highly skewed toward males is the cause of extinction because there are no more females to produce offspring (Figure 1B). A few extra generations easily make up for an increase of an order of magnitude in population size. *Conclusion 1: population decline due to a driving sex chromosome is approximately logistic, meaning the expected time to extinction increases approximately linearly with the log of population size.* As will be shown below, conclusion 1 applies for X drive as well.

Our goal is to determine the probability that a population can save itself before going extinct, so we would like to know the total expected number of suppressor mutations that will arise in the population of X chromosomes before the population goes extinct. The number of X chromosomes at generation *t* is *X*(*t*) = 3*XX*(*t*) + *XY*^*SR*^ (*t*), where *XX*(*t*) denotes the number of females at time *t*, *XY*^*SR*^(*t*) denotes the number of sex-ratio males at time *t*, and the factor of three accounts for the two X chromosomes found in females and the one found in XY males (found at the same frequency as females in this special case). Following logic similar to that for equation (1), the number of X chromosomes at generation *t* (Figure 1B) is

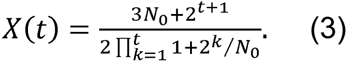

With mutation rate *μ*, the expected number of suppressing mutants is *X*(*t*)*μ* and the total number of mutants expected is

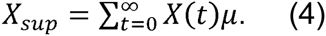

Without an analytical solution for the time at which extinction occurs, we sum to infinity because once the population size is near zero, the number of new mutations is negligible. This means only about 31,600 X chromosomes exist before extinction if the starting population is 2000, whereas almost 30 million exist before extinction with a starting population of one million. Nevertheless, even with a large starting population size, the mutation rate to a suppressor must be quite high to provide any chance of saving a relatively small population.

The selective advantage of a suppressor depends on the background on which it finds itself, which is dependent on the frequency of the driver in the population. Even without fitness costs, the suppressor only provides a fitness benefit when found in males carrying the driver (not in females and in non-driving males where it is neutral). The absolute fitness of a suppressing X chromosome (one that completely restores Fisherian sex ratios and has no fitness effects) in a driving male (XY^SR^) is (1 – *d*_*s*_)*R*_0_ so the total absolute fitness of a suppressing X at time *t* is

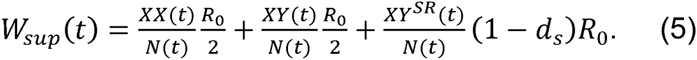

Assuming *R*_*0*_ =2 (as in Hamilton’s model) and *d*_*s*_=0.5 (complete suppression of drive), equation (8) becomes (*XX*(*t*) + *XY*(*t*) + *XY*^*SR*^(*t*))/*N*(*t*) = 1. This is a problem for the Hamilton scenario, because it means that even if a suppressor arises and takes hold, the population cannot deterministically increase in size from the size when it arose. Therefore, even populations with suppressors of Y drive are at risk of extinction due to stochastic variation around this new equilibrium population size. Though Hamilton’s model is obviously not realistic, the preceding suggests that populations with relatively low lifetime fecundity may be at higher risk of extinction than those with higher fecundity.

*Conclusion 2*: *the fate of a population with a driving sex chromosome is dependent on the relationship between the strength of drive and female fecundity. Species with low lifetime fecundity will be driven extinct by even weak drive (barring suppression).*

Suppressors may arise on three different genetic backgrounds: XX females, XY males and XY^SR^ males. In the background of females and standard males, the suppressor provides no benefit and must survive drift (at least two generations when a suppressor arises in XY males) before it has any selective advantage. In contrast, a suppressing X in SR males has a selective advantage right away assuming that male has the opportunity to mate. These differences provide us with some insight into why Y drive populations are so likely go extinct. Suppressors that arise early, when drivers are rare, are likely to find themselves in a non-driving background and therefore will be lost due to drift before they can spread. When X chromosomes are most common, and therefore the mutational target size is largest, there is little advantage to carrying a suppressor. Conversely, when the selective advantage of a suppressor becomes large relative to wild type (and it becomes very large), there are many fewer X chromosome mutational targets left. Furthermore, when the XY^SR^ genotype is prominent, since the sex ratio is so skewed toward males, a male that has both the suppressor and the driving Y (*X*^*sup*^*Y*^*SR*^) is unlikely to find a mate! *Conclusion 3: a) When the driver is rare, a rare suppressor is unlikely to find itself paired with a driver and will likely be lost due to drift; b) when the driver is common, the number of mutational targets (opposite sex chromosome) is small, so new mutations are unlikely; c) for Y drive, when the driver is common most males will go unmated since there is a scarcity of females so even X^sup^Y^SR^ males may not produce any daughters.* As discussed below, conclusions 3a and 3b apply to X drive, but for X drive, 3c is the opposite, when the driver is common, there is a scarcity of males, so the suppressor rapidly invades because any X^SR^Y^sup^ male is likely to mate (multiply) and produce sons.

#### Expansion of Hamilton’s Model for X drive

The case of a driving X chromosome is initially more difficult because there are now five possible genotypes (even before a suppressor arises). Recursions for the case of a driving X under Hamilton’s scenario are given in Supplemental Text 2 (Equation SB.5), however, no further analytical solutions will be presented. Numerical solutions are, however, both straightforward and useful (Figure 1C and 1D).

Again, the time to extinction is approximately linearly related to the log of the starting population size (*Conclusion 1* above): with times to extinction of 46, 57 and 67 generations for populations of 10^4^, 10^5^ and 10^6^ respectively. As seen in Figure 1D, the sex-ratio and number of mutation targets (Y chromosomes in this case) behave similarly to the Y drive model, except that the process takes longer and the sex-ratio goes in the opposite direction. For comparison, about 25,000 Y chromosomes exist before extinction with a population of 2000, but 25 million exist with a starting population of 1,000,000. Even with the longer time to extinction with X drive, the number of Y chromosomes that exist while a driving X leads to extinction is less than the number of X chromosomes during the advance of a driving Y, because the when sex ratios are equal, there are three times as many X chromosomes as Y.

### Analytical progress beyond Hamilton’s Model

Relaxing the assumptions of Hamilton’s model significantly muddies the waters since there are so many potential parameters involved (Table 1). The parameters can be divided into those involved in demographics (*N*_0_ and *μ*), reproduction (*M* and *R*_0_), drive (*d* and *d*_*s*_) and fitness (all selection coefficients - *s*). For simplicity, we begin by assuming that the drive allele carries no fitness costs in either sex. This seems reasonable since fitness costs are thought to be associated with deleterious mutations linked to the driving locus by inversions (Jaenike 2001; Burt and Trivers 2006). These inversions also link enhancers of drive and therefore are selectively favored. As a new drive locus begins to spread, however, it is unlikely that such an inversion will have time to arise. We acknowledge that the drive locus may also have direct fitness costs, but ignore this possibility for now. We also assume that suppressors of drive are perfect so that *d*_*s*_ is always 0.5 and males carrying the driver and suppressor have 50% daughters. We have already dealt with population size (*N*_*0*_) and mutation rate (*μ*), leaving three parameters (*M, R*_0_ and *d*) to explore.

#### Y drive – When is extinction deterministic?

The first concern is under what circumstances extinction is certain (barring a mutation that suppresses drive and given that the driver actually invades). Much theoretical work has shown that stable equilibria exist for many parameter sets (Clark 1987; Carvalho and Vaz 1999; Jaenike 2001; Hall 2004), this is supported up by the occurrence of seemingly stable equilibria in natural populations (reviewed in Jaenike 2001). This does not suggest, however, that all drivers reach a stable equilibrium, and indeed there is a strong ascertainment bias to only sample populations not on their way to immediate extinction. There are two ways in which a population may go extinct due to a driving sex chromosome. First, the sex that is driven against may be removed from the population – the equilibrium number of individuals of that sex is zero or less than one. Second, the sex that is driven against may reach an equilibrium frequency, but the absolute number may be small enough that the sex is likely to be lost due to drift. For example, if the equilibrium number of females in a population is 10, there is a good chance that in one generation zero females will be produced and the population is therefore doomed.

For the driving Y, the situation is quite simple. A population will be driven to extinction by the driving Y if the average number of females produced is less than one. Since mothers produce *R*_*0*_ offspring and *R*_*0*_(1-*d*) of them will be female, for populations to survive

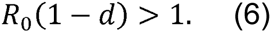

Most cases of sex-ratio drive that have been studied show drive that is strong (between 0.9 and 1.0), although there is surely an ascertainment bias as weak drivers are less likely to be noticed. The average number of offspring produced per mother is extremely variable. With strong drive (*d*>0.9), *R*_0_ would need to be greater than ten for populations to have any chance of survival.

#### Y drive – when is extinction risk stochastic?

Of course, if *R*_*0*_(1-*d*) is greater than one, but still small, the population may go extinct due to drift. The equilibrium number of females is *N*_*0*_(1-*d*), so extinction is likely if this number is small. Therefore, even intermediate population sizes (*N*_*0*_=10,000) and very strong drive (*d*=0.99), the population may go extinct since at fixation of the driving Y, the expected number of females is 100. For weak drive, populations might go extinct if the driver invades, but the driver’s chance of invading is quite low. For example, for *R*_*0*_=2 and *d*=0.6, the inequality in Equation 6 is not satisfied, but with such weak drive and only two offspring per female, the driver almost never invades to begin with (simulations show that the driver is lost almost 90% of the time).

#### Y drive: How long to extinction?

Recursion equations for a driving Y with additional parameters discussed above are given in Supplemental Text 2. Now, however, the population size may become larger than the carrying capacity, so if *XX*′ + *XY*′ + *XY*^*SR*′^ = *N*′ >*N*_0_, we must reduce each genotype proportionally so that *N*′ = *N*_0_. Even in the case of unregulated population growth, the time to extinction is only affected slightly, even when the population gets extremely large. For example, with **N**=2000, *P*=10, perfect drive and no fitness cost of the driver, the maximum population reached without regulation is in excess of 100 billion, the population goes extinct in 23 generations as opposed to 19 generations in the same population with regulation. The reason for this is obvious – once females are limiting, the population crashes extremely fast no matter what its size. The effect of population regulation on extinction time grows rapidly with *d*<1.0.

Interestingly, since there is no fitness cost to females of a driving Y (since the Y never occurs in females) and since the Y experiences haploid transmission, there is no stable equilibrium for Y drive, assuming no effect of sperm competition.

*Conclusion 4: Unlike populations with X drive, those with Y drive do not reach a polymorphic equilibrium for the driving Y. The Y either fixes or is lost.* This may provide another reason that Y drive is rarely reported in natural populations. Note that Clark (1987) found limited parameter space allowing for the maintenance of a driving Y chromosome, but this was in the presence of a driving X chromosome which we do not allow in the current scenario.

For reasons described above in addition to this density dependent control, further analytical solutions were not attempted. Figure 2A shows the time to extinction (from simulation over a wide range of parameter values). Note that as drive becomes stronger, the time to extinction is less affected by the number of offspring produced by females in their lifetime. The figure again confirms that starting population size has little effect on time to extinction.

**Figure 2:**
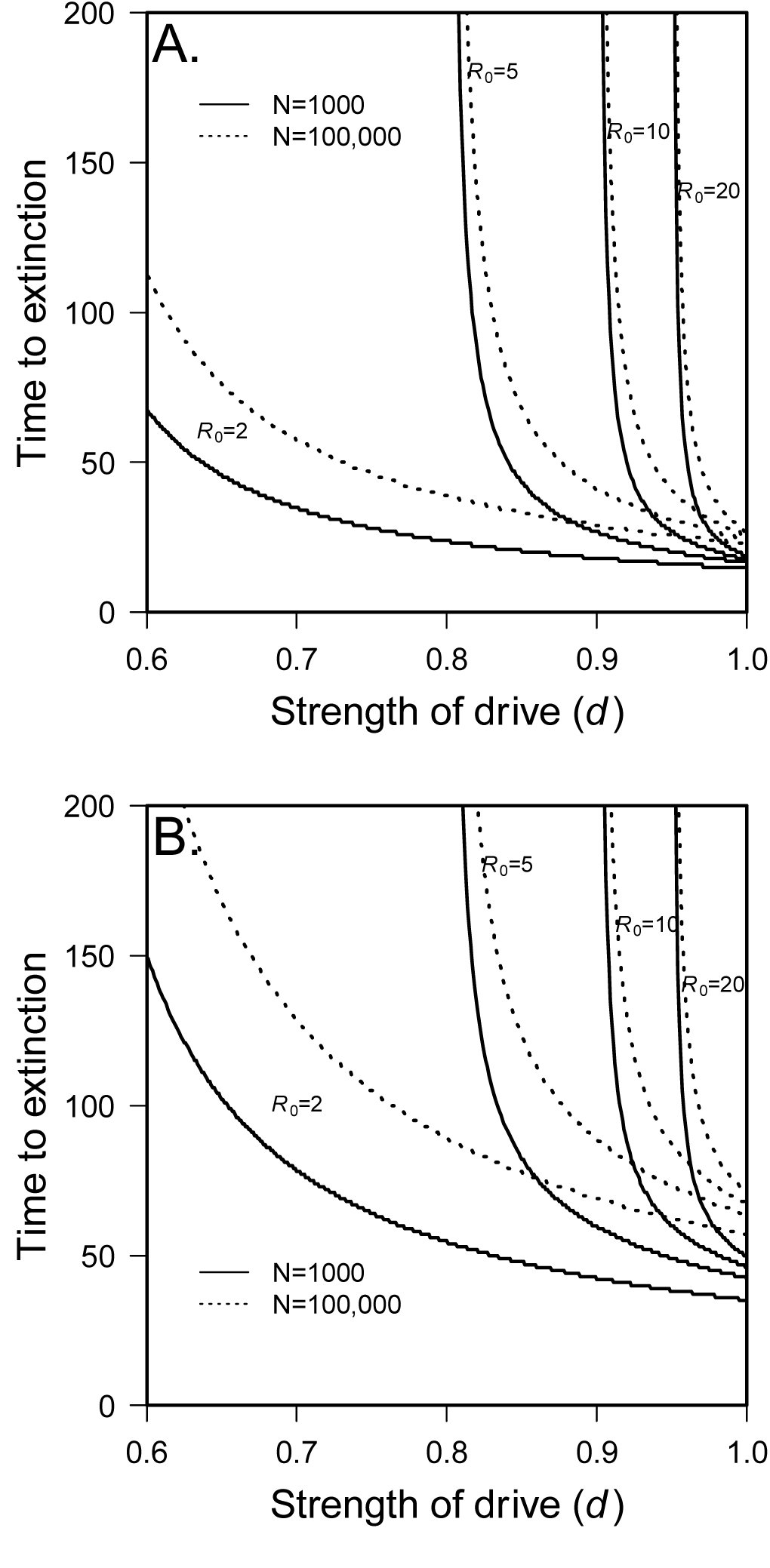
The deterministic time to extinction is strongly influenced by strength of drive (without evolution of suppressors) A) Y drive B) X drive. Starting population size: N_0_=1000 (dashed line); N_0_=1000000 (solid line), other parameters as described by Hamilton.

#### X drive and extinction

With a driving X chromosome, the population will go extinct if females do not produce on average at least one son. Assuming no fitness costs of the driver (in either sex), the same inequality applies as for Y drive (Equation 6). With fitness costs of the driver in females, an equilibrium frequency of the driver may exist and prevent extinction. The same applies for stochastic extinction. If *N*_*0*_(1-*d*) is small enough, populations may contain so few males that they go extinct due to a chance loss of males. This suggests another reason that populations with Y drive are more likely to be doomed: anisogamy.

*Conclusion 5: Assuming male remating rate is higher than female remating rate, males are more easily able to compensate for a population level sperm shortage due to a scarcity of males than females are able to compensate for a population level egg shortage due to a scarcity of females.*

#### X drive: How long to extinction?

Figure 2B shows the time to extinction (from simulation over a wide range of parameter values). Though the conditions for extinction are the same for X and Y drive, extinction takes much longer for X drive. This is true for two reasons: 1) there are three times as many X chromosomes as Y, so a sweep takes longer and 2) as Hamilton (1967) noted, while Y drive occurs every generation, a driving X only finds itself in males one-third of the time and can only drive in alternate generations.

### Simulation results

#### Hamilton’s parameters

Using Hamilton’s parameters and simulations allowing for stochasticity, populations with a driving Y chromosome are doomed unless the mutation rate is very high (*e.g*. a mutation rate of 10^-3^ yields a probability of 0.06) or standing genetic variation contains several copies of the suppressing X chromosome (Figure 3A). The prognosis for populations with X drive is much better (Figure 3B). For new mutation, the probability of survival is 0.07 for a mutation rate of 10^-5^, so similar probability of population survival occur at a mutation rate two orders of magnitude lower than for Y drive. Note, however, that 10^-5^ is probably quite high for a mutation rate. Populations with a driving X can be saved by standing genetic variation with a relatively small number of mutants. Ten starting mutants are required for a 0.07 chance of survival in Y drive, but only 1 starting mutant is required for the same probability with X drive. While the mutation rates discussed above may seem exceedingly high, it is important to note that suppression of drive may involve the loss of repetitive satellite DNA which may occur at rates much higher than the nucleotide mutation rate. Hamilton’s parameters are obviously biologically unrealistic, but they support two important ideas. First, as described above, populations are at a much greater risk of extinction caused by a driving Y than a driving X. Second, with plausible parameter values, standing genetic variation is much more likely to save the population than new mutation (see also Orr and Unckless 2008).

**Figure 3:**
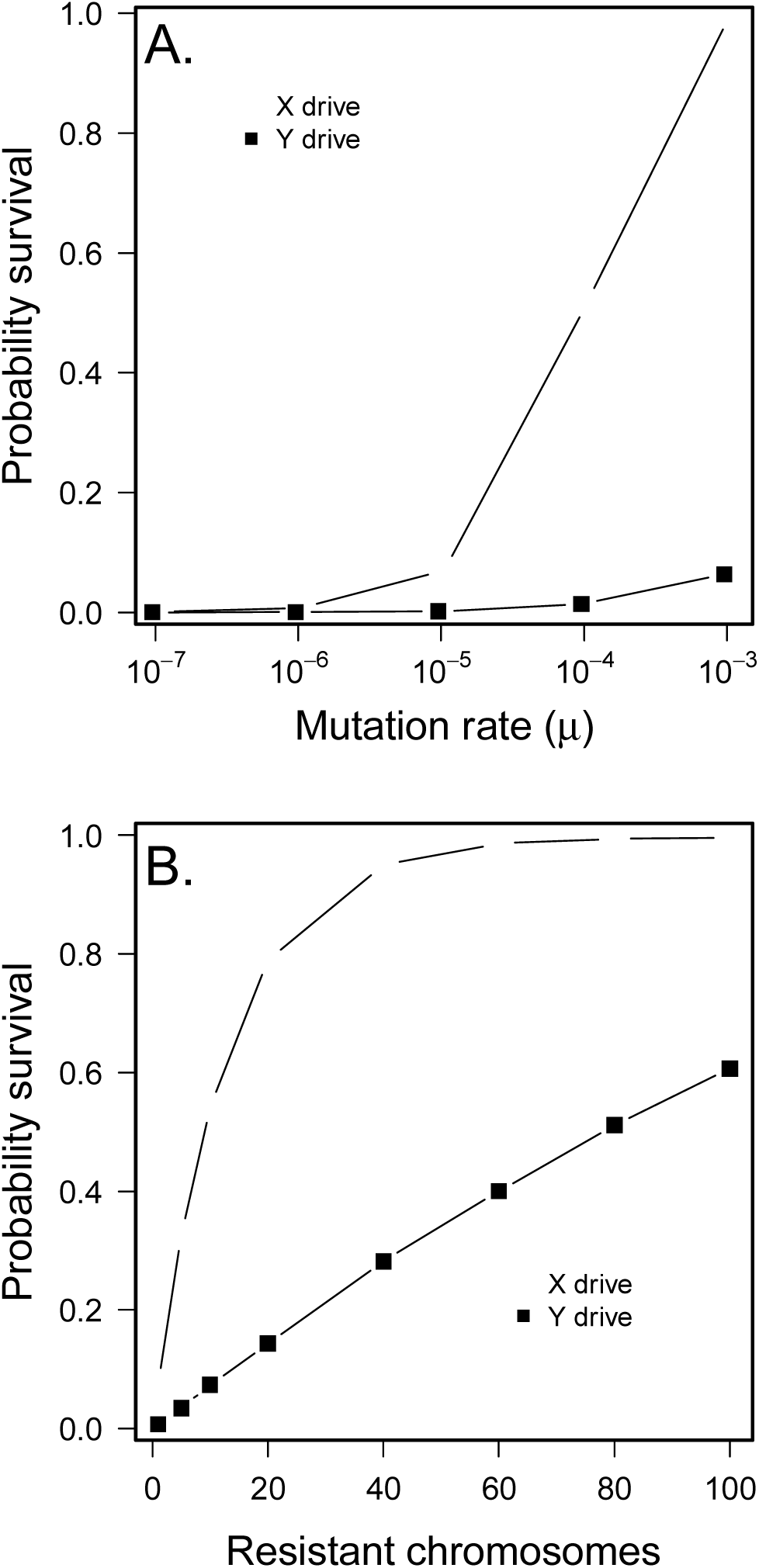
Probability of survival with Hamilton’s parameters is much greater for X drive than Y drive, A) new mutation and B) standing genetic variation for resistant chromosomes. Data from 100,000 simulation realizations.

#### Beyond Hamilton’s parameters

We now examine the fate of populations when parameter values are more realistic. It would seem that Hamilton’s parameters (small population size, two offspring per female, perfect drive, etc.) represent the dire extreme. Increasing any of the parameters in Table 1 (other than fitness costs of the suppressor) should increase the probability that populations survive. The difficulty in exploring parameter space is that with nine possible parameters, even with only two values for each parameter, there are 512 parameter sets. We therefore focus on those that might be most relevant during the initial spread of a driver – *N*_0_, *P* and *d*. Then we consider each of the fitness effects individually, discussing interesting interactions between parameters as appropriate.

Results of simulations for both X and Y drive and from new mutation and standing genetic variation are shown in Figure 4. The mutation rate to suppressors for Y drive is 10^-4^, and for X driver it is 10^-6^ to achieve an intermediate level of population survival for both scenarios.

**Figure 4:**
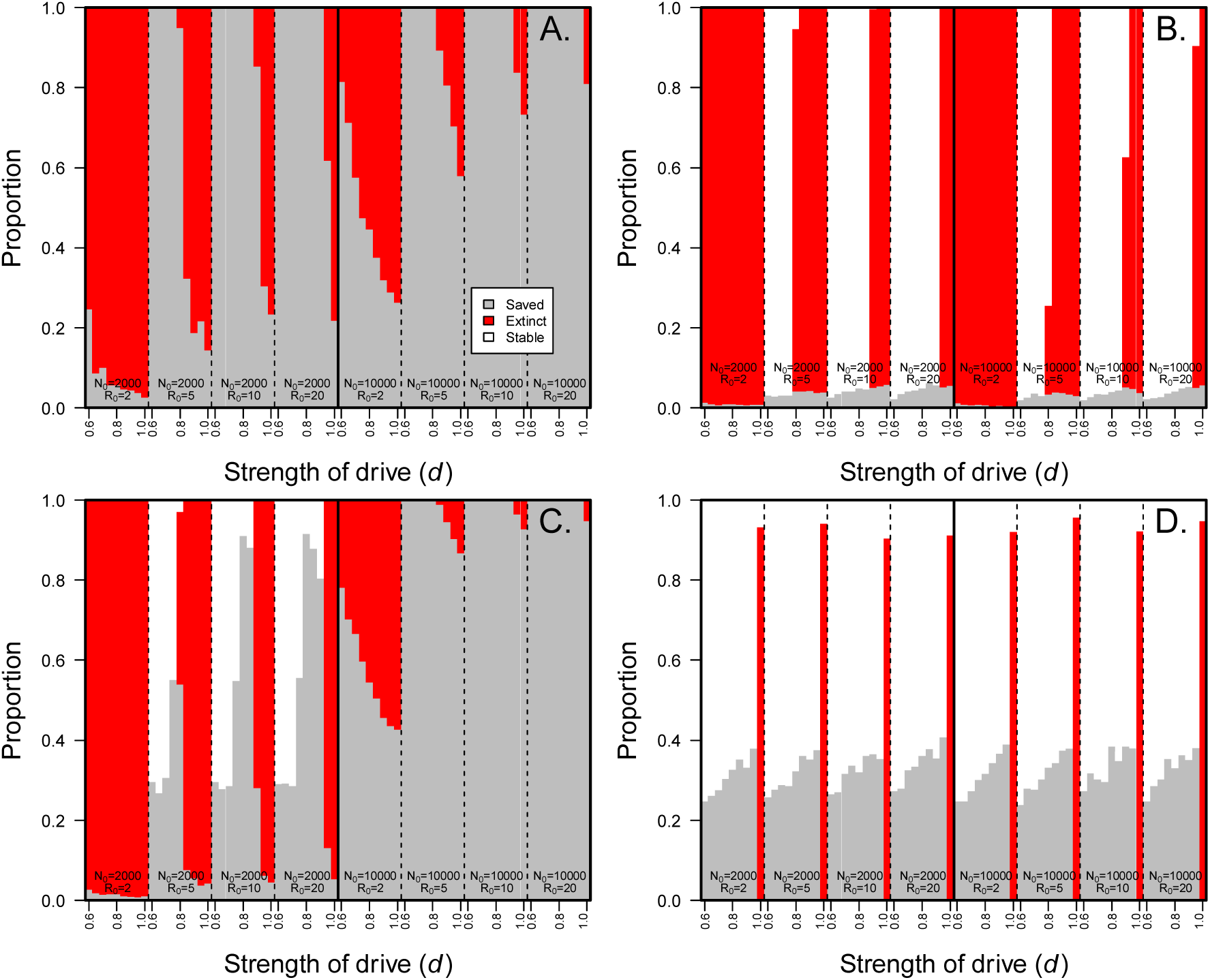
Fate of populations with driving chromosomes with no fitness costs of driver or suppressor A) Y drive from new mutation (u=10^-4^), B) Y drive from standing variation (1 mutant at time t=0), C) X drive from new mutation (u=10^-6^), D) X drive from standing variation (1 mutant at time t=0). Only simulations where the driver successfully invades are presented.

For Y drive from new mutation, results are as intuitively expected (Figure 4A). First, The probability that the driver is lost decreases with increasing strength of drive (noted in red). Neither the initial population size (*N*_*0*_) nor reproductive rate (*R*_*0*_) have much influence on the probability that a driver invades. This is true because what is critical for the driver is what happens in the first few generations. During this time the population size is close to carrying capacity, so females produce an average of two offspring surviving to reproductive age regardless of their number of births. The actual carrying capacity also has very little to do with the invasion of a driver. The strength of drive, on the other hand, determines whether the expected number of sons (and therefore driving chromosomes in the next generation) is 2 (*d*=1) or 1.2 (*d*=0.6). This has a profound effect on a driver’s ability to invade. Figure 4B confirms that the ability of a driver to invade has little to do with whether or not suppressors come from standing variation or new mutation (as long as the frequency of suppressors is low). As discussed above, without the existence of a driving X, Y drive does not lead to stable equilibrium. Either the driver fixes or it is lost. The sex-ratio can reach a stable equilibrium assuming fixation of the driver if the strength of drive is less than one. For Y drive with suppression from new mutation, since new mutation can occur at any time, all populations either were saved by a suppressor or went extinct (Figure 4A). As expected, the probability that a suppressor saves the population decreases with the strength of drive, increases with *R*_*0*_ and increases with population size. Note, that while figure 4A suggest that large populations with large reproductive capacities (most insects) are at low risk of extinction even with strong drive, the mutation rate to suppressors (10^-4^) is very high in this case. Simulations show that with an initial population size of one million, *R*_*0*_=20, *μ*=10^-7^, and *d*=1, given the driver invaded, the population survived only about 18 percent of the time.

The situation is similar for Y drive and suppressors from standing variation (Figure 4B), except now if the suppressor is lost due to drift, the population can become stable *without* the invasion of the suppressor as long as *R*_*0*_>2. Figure 4B also shows under what circumstances drift might drive a population extinct. This is so whenever under any particular parameter set, both extinction and stability were found. For example, with *N*_*0*_=2000, *R*_*0*_=5 and *d*=0.8, the equilibrium number of females in the population is 400, but extinction occurs about three percent of the time. Figure 4B also shows that with a single suppressor in the population at time *t*=0, the population size has little influence on whether or not a suppressor saves the population. Population size does, however, influence whether the population will remain stable or go extinct.

Again, the prognosis for populations with X drive is much better. In fact, we have used a base mutation rate of 10^-6^ instead of 10^-5^ as in Y drive. This makes comparison of X and Y more difficult, but using 10^-4^ would nearly always result in populations with X drive being saved by new mutation. Figure 4C shows the results of simulations for X drive with the same parameters as for Y drive (except mutation rate). Note that under Hamilton’s parameters, the population is almost certainly doomed, but with greater *R*_*0*_ or weaker drive, the population is likely to reach a stable equilibrium or be saved by a suppressor. With 10,000 as a starting population size, the population almost never goes extinct with 10^-6^ as a mutation rate. Interestingly, the probability that a suppressor invades spikes as *d* approaches a value where extinction is deterministic. This is because selection for a suppressor is greater the stronger the drive. If drive is too strong, however, extinction is a possibility and the probability that a suppressor invades is lower because the population goes extinct before the suppressor can invade. The situation from standing variation is qualitatively similar, but since a suppressor lost by chance cannot be regained, suppressors are less likely to gain ground with weaker drive. *Conclusion 6: Suppressors are most likely to become established when the strength of drive is great enough that there is strong selection for suppression, but weak enough that the population does not go extinct too quickly.*

#### Fitness consequences of drivers and suppressors

Empirical work on sex-ratio and autosomal drive show that the driver and suppressor often have significant fitness consequences to their hosts (Jaenike 2001). In these cases it is not known whether this is due to direct costs of the driver or suppressor or linked deleterious mutations that accumulate as the driver/suppressor sweeps. When considering extinction, linked deleterious mutations are less likely to be an issue because given how fast strong drivers sweep through a population, there is unlikely enough time for several deleterious mutations to accumulate. If the driver locus itself carries some fitness cost in either sex, this would presumably slow down its spread and reduce the risk of extinction. Conversely, costs of a suppressor should reduce its chances of invading, and therefore increase the chance of extinction. Again, a population with a driving Y is at a disadvantage because a suppressing X is only in males one-third of the time – more as the sex ratio becomes more skewed. So, any cost in females will reduce its chances of spreading. A Y chromosome that suppresses X drive is *always* in males and therefore carries no fitness cost to females. Of course, autosomal suppressors are equally likely in both sexes when sex-ratios are balanced. So they may do better combating Y drive since the sex ratio skews toward males, the sex in which they can actually suppress the drive.

Tables S1 and S2 show the results when fitness parameters are allowed to vary. We kept fitness costs relatively low (0.1) because this might be realistic biologically if costs are direct and not due to linked deleterious mutations. We also include a dominance coefficient (*h*) whenever the driver or suppressor finds itself in the diploid state.

Fitness costs of the driver generally have small but real effects on the probability that a population escapes extinction. With restrictive parameter sets similar to those employed by Hamilton, moderate fitness costs have little influence on extinction risk because extinction is so quick regardless of associated driver costs and in the few cases where a suppressor escapes stochastic loss, selection for suppression well outweighs the associated suppressor costs. With a higher reproductive output, fitness costs have a moderate impact on population survival with no cost of the driver. If drive is weaker *(d*=0.9), populations are less likely to go extinct) however a striking patterns still exists. Those populations threatened by extinction are unlikely to be saved regardless of fitness costs, at least with the parameters used.

#### Given population survival, how was it saved?

In those populations that were saved by suppressors from new mutation, those mutations arose early on. The mean time until a successful suppressor of Y drive arose was usually between 20 and 30 percent of the mean time to extinction, and given appreciable extinction risk, this was within the first ten generations after driver introduction. For example, with Hamilton’s parameters and a mutation rate of 10^-4^, the mean time until a successful suppressor arose was about 3.6 generations, while the mean time to extinction was about 15 generations. The time until a successful suppressor arose decreased asymptotically with increasing strength of drive (Figure 5A). The same patterns were true for successful suppressors of X drive. With Hamilton’s parameters and a mutation rate of 10^-5^, the mean time was about six generations, but the time to extinction was much longer. Figure 5B shows the time until a successful suppressor arose with increasing strength of X drive. Patterns are broadly similar for X and Y drive, but X drivers tend to allow for more time until successful suppressors arise since the time to extinction is also longer.

**Figure 5:**
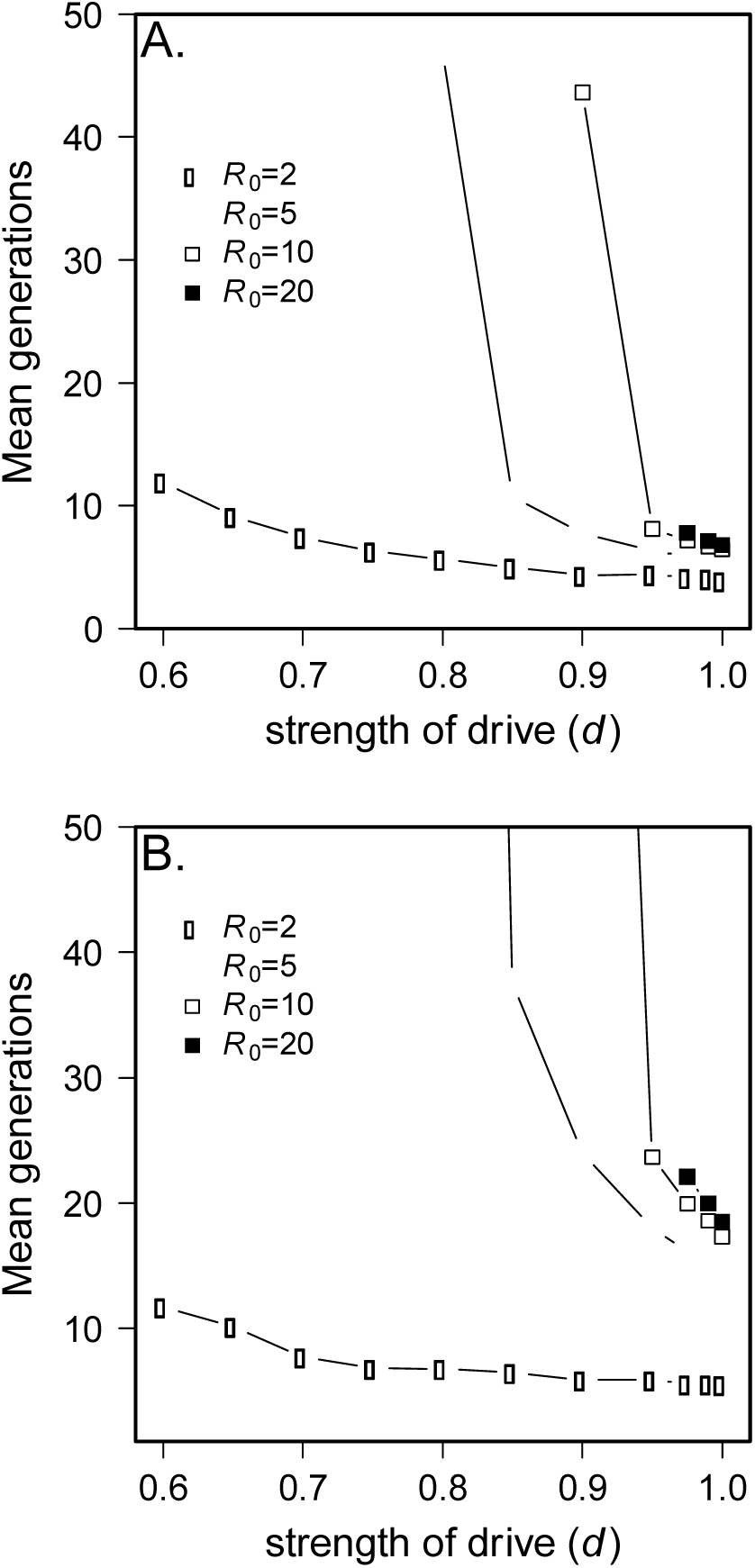
Mean time until a successful suppressor arises (given that it does) A) X drive with *N*_*0*_=2000, *u*=10^-5^, *M*=1, and no fitness costs, B) Y drive with *N*_*0*_=2000, *u*=10^-4^, *M*=2 and no fitness costs. Only parameter combinations where extinction was deterministic (barring suppression) are plotted. All realizations assume perfect suppression (*d*_*s*_=0.5), 10,000 realizations per parameter set. In panel B cutoff values are 274.5 generations for *d*=0.8 and *R*_*0*_=5 and 154.6 generations for *d*=0.9 and *R*_*0*_=10.

### Discussion

Although sex-ratio meiotic drive is widespread among sexual species (Jaenike 2001) and the implications for extinction caused by drive have been recognized for quite some time (Hamilton 1967), few studies have examined the consequences of drive on populations. In this paper, the link between sex-ratio meiotic drive, its suppression, and extinction was modeled explicitly, and simulations were employed to begin to explore the parameter space. We find that population size has only a small influence on the time to extinction which scales with the log of population size (*Conclusion 1*). For several reasons, Y-linked drivers, should they arise, are likely to drive populations to extinction. X-linked drivers, on the other hand, pose less of a risk of extinction.

Despite efforts to examine biologically reasonable parameter values, the simulations showed that extinction *was* likely after sex chromosome drivers are introduced. There are practical and biological reasons for this. First, simulations that either take many generations for extinction or are likely to lead to a stable equilibrium take much longer to run. There is considerable variation in life history parameters associated with the several groups of organisms exhibiting drive. Furthermore, even though *Drosophila* are capable of producing hundreds of offspring throughout their lifetime, their actual fecundity in the wild is surely much lower. Rosewell and Shorrocks and references therein (1987) estimated daily survival in several *Drosophila* species ranging from 0.42 to 0.83. Assuming that mortality is geometrically distributed, the mean survival of a given fly is the reciprocal of its daily mortality. So, even if daily mortality is about 1/3 (in lines with estimates for *D. melanogaster*), flies only survive three days on average. Assuming daily fecundity is about 60 eggs and constant and that flies don’t lay eggs for the first two days (Novoseltsev et al. 2005), the average lifetime fecundity is about 50. Of course, the variance is huge, scaling with the square of fecundity. Finally, *Conclusion 2* suggests that one reason we might see drive more often in species with high lifetime fecundity is that extinction is much more likely with lower fecundity.

Most theoretical studies of sex-ratio (and autosomal) drive have (rightfully) used empirical estimates of drive parameters from the populations with balanced drivers. This may have the effect of misleading us from two directions. First, weak sex-ratio drivers (*d*≈0.55) could sweep through a population without notice since such minor deviations from equal sex-ratios would only be detected if under careful scrutiny (Corbett-Detig and Hartl 2012). Suppressors could also restore normal sex-ratios without notice. As, discussed by Hartl (1970), even weak drivers and their suppressors are under very strong selection. Second, very strong drivers, especially those without fitness consequences, are likely to go undetected if they lead to population extinction. Thus the empirical values described in the literature may not be appropriate for the extinction scenario.

Driving Y chromosomes appear to be much more rare than driving X chromosomes (Jaenike 2001). There are several possible reasons for this. First, a degraded Y chromosome may lack sufficient starting genetic material for drive to arise *de novo.* In fact, the only well-characterized case of Y drive is found in *Aedes aegypti* in which the X and Y are homomorphic (Mori et al. 2004). Second, driving Y chromosomes are more likely to lead to population extinction. As noted by Hamilton (Hamilton 1967), a driving Y is *always* in males, so unlike the X chromosome which spends at least 2/3 of its time in females, the Y can drive every generation. Because of this, with similar parameters, Y drivers lead to extinction in less than half as many generations as X drivers. Similarly, if it is assumed that the easiest way to suppress drive is to mutate at the responder locus, an X-linked suppressor only finds itself in males one-third of the time (though this will increase as the sex-ratio becomes skewed toward males), but a Y-linked suppressor is found only in males. Perhaps most damning for populations with Y drive is that when the driver is common and the sex-ratio is skewed toward males, most males will go unmated, so even if a suppressor arises in a male, it will very likely be lost because the male fails to secure any matings (*Conclusion 3* above). Even if an X-linked suppressor arises in a female, to save the population, it must be passed to male offspring, and those male offspring must be lucky enough to mate. The scenario is the opposite for X drive. As the driver becomes common and the sex-ratio becomes skewed toward females, males are limiting and therefore males with a suppressing Y are likely to mate multiple times. We also don’t expect to find Y drivers segregating at intermediate frequencies since, unlike X drive, the fitness advantage for a driving Y in males cannot be balanced by costs in females (*Conclusion 4*). Finally, anisogamy may also contribute to the lack of empirical examples of Y drive since populations are likely to be able to cope with a shortage of sperm more easily than a shortage of eggs (*Conclusion 5*).

If suppressors are allowed to occur on autosomes some of the results presented above may indeed be different. First, with both types of drive, the number of autosomes is always larger than the number of opposite sex chromosome so there are more mutational targets. On the other hand, mutation rates to autosomal suppressors may be several orders of magnitude less than those on the opposite sex chromosome, since evolving insensitivity at the responder locus may be as simple as altering the number of tandem repeats at that locus. An active suppressor on the autosome is presumably much harder to evolve. Since both sexes have two copies of autosomal genes, the relative proportion of the time an autosome finds itself in one sex is equal to the sex-ratio at that time. Autosomal suppressors of Y drive will spend more time in males (where they can actually suppress drive) than in females because as the driver spreads, the sex-ratio becomes male-biased. The opposite is true for X drive – as the X driver spreads, the autosomes are more likely to be found in females since the sex-ratio is female biased.

The models presented above incorporate a fitness cost of the driver in males, but this is different from sperm competition models analyzed by Taylor and Jaenike (2002; 2003) because fitness costs in their model were frequency dependent: a male carrying a driving X suffers no fertility cost after a single mating, but when mating multiply, fertility is reduced. As the sex-ratio becomes skewed toward females, males are more likely to mate multiply and fertility costs of drive are realized, especially when females have mated with both standard (ST) males and those carrying the driver (SR). This creates an unstable equilibrium, above which males are so limiting that females are likely to mate with only one male and therefore no sperm competition occurs. Furthermore, with high frequency of the driver, the relative fitness cost of drive is reduced since most males carry the driver. Interestingly, Y drive *may* lead to a stable equilibrium with sperm competition because as the sex-ratio becomes skewed toward males, *all* females are likely to be multiply mated meaning that sperm from males carrying the driver are likely to suffer from sperm competition.

Our results provide important implications for pest control strategies and conservation. If the goal is to eliminate a pest population completely, the introduction of a strong Y driver is likely the best strategy. Knocking down the total population size in addition to the introduction of a driver would be useful, not because it will have a large effect on the time to extinction, but because it would decrease the chances that a suppressor might be segregating in the standing genetic variation.

Sex-ratio meiotic drive has impacted a wide variety of evolutionary and ecological processes, including interspecific competition (James and Jaenike 1990; Unckless and Clark 2014), reproductive isolation (reviewed in McDermott and Noor 2010), changes in mating systems (Price et al. 2008; Pinzone and Dyer 2013), sex-chromosome rearrangement and sex determination (Kozielska et al. 2010), the maintenance of genetic variation (Carvalho et al. 1997; Jaenike 1999; Montchamp-Moreau et al. 2001; Branco et al. 2013; Unckless et al. 2015) and population extinction (Hamilton 1967 and this paper). Given the potency of meiotic drive to influence so many ecological and evolutionary processes, its role as an important architect of evolutionary change is now clear. The universality of this role depends on the incidence of drive across taxa. While we know that meiotic drive exists in plants, mammals, insects and nematodes, examples tend to come from well-studied systems so the proportion of species affected by drive is still unknown.

